# Protein substrates engage the lumen of O-GlcNac transferase’s tetratricopeptide repeat domain in different ways

**DOI:** 10.1101/2020.12.22.423982

**Authors:** Cassandra M. Joiner, Forrest A. Hammel, John Janetzko, Suzanne Walker

## Abstract

Glycosylation of nuclear and cytoplasmic proteins is an essential post-translational modification in mammals. O-GlcNAc transferase (OGT), the sole enzyme responsible for this modification, glycosylates over a thousand unique nuclear and cytoplasmic substrates. How OGT selects its substrates is a fundamental question that must be answered to understand OGT’s unusual biology. OGT contains a long tetratricopeptide repeat (TPR) domain that has been implicated in substrate selection, but there is almost no information about how changes to this domain affect glycosylation of individual substrates. Here, we used proteome-wide glycosylation profiling and probed glycosylation of selected purified substrates to show that asparagine and aspartate ladders that extend the full length of OGT’s TPR lumen control substrate glycosylation. We also found that substrates with glycosylation sites close to the C-terminus bypass lumenal binding. Our findings demonstrate that substrates can engage OGT in a variety of different ways for glycosylation.

## TEXT

O-linked β-N-acetylglucosamine (O-GlcNAc) transferase (OGT) is an essential nutrient- and stress-responsive enzyme that transfers GlcNAc to serine or threonine side chains of thousands of nuclear and cytoplasmic proteins.^*1-6*^ OGT’s substrates are structurally and functionally diverse and are involved in most cellular processes.^*7-15*^ A longstanding question in the field is how OGT chooses its substrates. OGT contains a C-terminal glycosyltransferase domain that performs the chemistry and an N-terminal tetratricopeptide repeat (TPR) domain that has been implicated in substrate selection (Figure 1).^*3, 16-22*^ OGT’s TPR domain, which contains 13.5 repeats, forms a superhelix with conserved asparagine and aspartate ladders that extend the length of the lumenal surface (Figure 1A & Tables S1-2).^*18, 23*^ A structure of OGT complexed with a peptide revealed a network of bidentate contacts from five sequential asparagine amides in the proximal TPR lumen to the peptide backbone; three aspartate sidechains contact polar side chains of the peptide (Figure 1).^*24*^ Proteome-wide studies showed that the active site-proximal asparagines and aspartates are important in substrate glycosylation, contributing to affinity and controlling selectivity, respectively.^*23, 25*^ These studies identified functions of residues in the proximal TPR lumen in substrate glycosylation, but the TPR lumen is more than 100 Å long and the functions of conserved asparagine and aspartate residues in the medial and distal lumen are not well understood. Elucidating the mechanisms of substrate selection is critical because elevated O-GlcNAc levels have been implicated in cancers and metabolic diseases.^*26-31*^ To evaluate opportunities for therapeutic intervention, we need information on how different regions of the OGT’s TPR domain are involved in glycosylation of different substrates.

**Figure 1.**
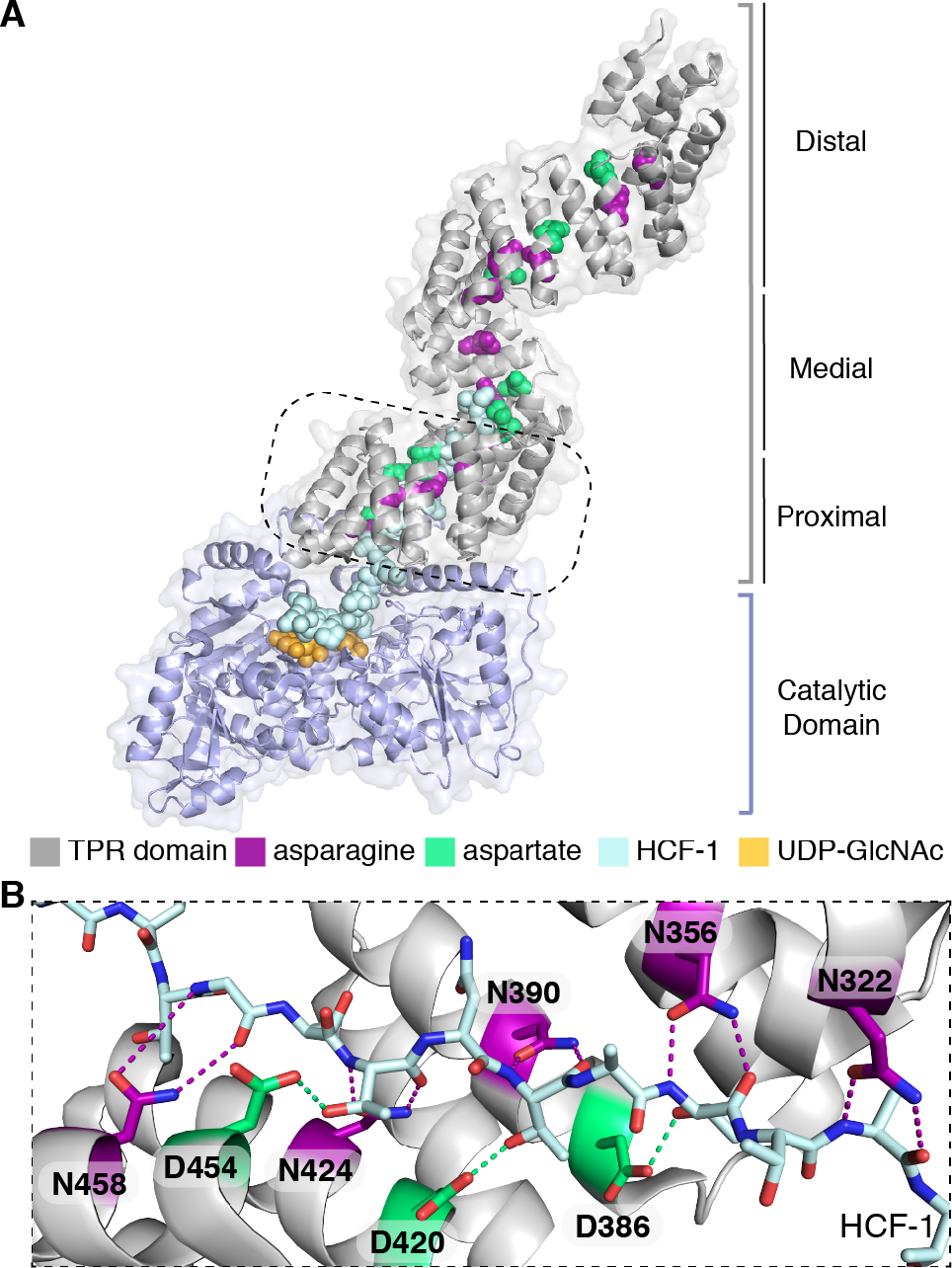
Conserved asparagine and aspartate ladders line the entire TPR lumen of OGT. (A) Composite structure of full-length OGT complexed with the HCF-1 substrate (light blue) shows active site and TPR lumen binding (PDB 1W3B and 4N3B). The TPR lumen is lined by conserved asparagine (magenta) and aspartate (green) ladder motifs. The proximal TPRs (dashed box) show contacts with the HCF-1 peptide. (B) Proximal asparagine and aspartate residues lining TPR lumen interact with HCF-1 peptide (90° turn view of dashed boxed in A).

To determine whether the entire asparagine ladder plays a role in glycosylation, we tested the proteome-wide activity of three OGT constructs in which groups of five asparagines were mutated to alanine (Figure 2A, referred to as N5A mutants). HeLa extracts were used for these studies because they are metabolically active and mimic a cellular environment, but enable precise control of the concentrations of OGT added.^*23, 25, 32*^ As reported previously, N5A_prox_ showed reduced global glycosylation compared to wildtype OGT.^*25*^ The N5A_med_ and N5A_dist_ mutants also showed lower glycosylation activity than wildtype, but glycosylation gradually increased as the asparagine mutations moved further from the active site (Figure 2B, see asterisks). Although the asparagines closest to the active site were more important, these results showed that the entire asparagine ladder contributes to global protein glycosylation.

**Figure 2.**
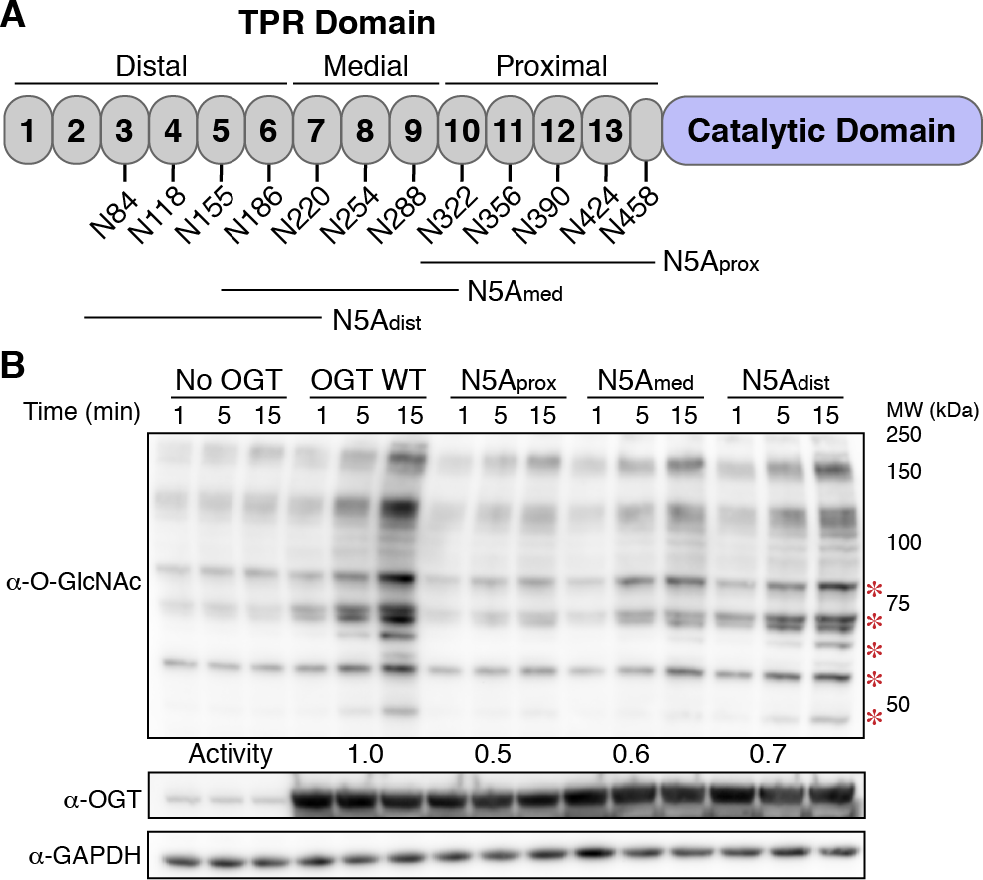
OGT requires the entire asparagine ladder for proteome-wide glycosylation. (A) Cartoon schematic of asparagine ladder mutations. (B) HeLa extracts shows loss of global glycosylation by OGT N5A mutants. Red asterisks highlight bands that are glycosylated at WT levels when most distal asparagines are mutated to alanine. Representative blot for two independent experiments. See methods for activity calculations.

We took a similar approach to assessing the roles of the aspartate ladder residues in global substrate selection. For these studies, we mutated conserved aspartates to alanines in groups of two (Figure 3A, referred to as D2A mutants). As previously reported, global glycosylation increased when aspartates in the proximal region of the TPR domain were mutated to alanines (D2A-2_prox_, Figures 3A & S1), highlighting the importance of these residues as “gatekeepers” that limit glycosylation activity.^*23*^ Global glycosylation also increased for other D2A mutants along the TPR lumen (Figure S1), with the greatest increase observed for mutations in the medial lumen (D2A-3_med_; Figure 3B). For different D2A constructs, we also observed pronounced differences in some product bands (compare D2A-2_prox_ to D2A-3_med_), suggesting that aspartates in different regions of the TPR lumen differentially regulate substrate glycosylation. When pairs of D2A mutants in the proximal and medial lumen were combined (D4A), bands observed for the individual pairs remained in the quadruple mutant (Figure 3B, see asterisks), suggesting that contributions of different aspartate pairs in controlling substrate selection are additive. Overall, the results for the aspartate ladder mutants implicate the entire TPR lumen in proteome-wide glycosylation, but show that the effects are largest for aspartates in the five TPRs closest to the active site (*i*.*e*., TPRs 9 to 13).

**Figure 3.**
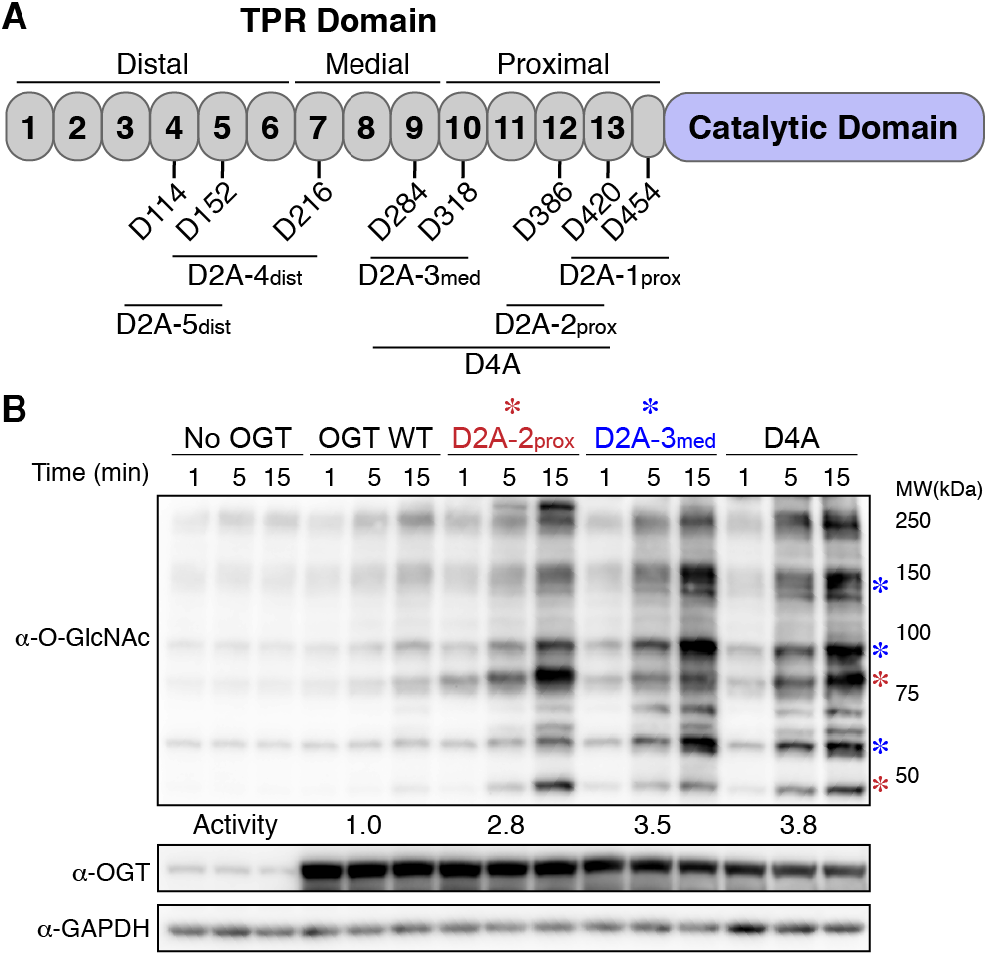
Aspartate ladder regulation of substrate selection is additive. (A) Cartoon schematic of aspartate ladder mutations. (B) Glycosylation of HeLa extracts by D2A-2_prox_ and D2A-3_med_ mutants show large global increase in glycosylation and preferential increases in different O-GlcNAc bands (see asterisks). These preferences are additive in the D4A mutant. Representative blot for two independent experiments. See methods for activity calculations.

The extract experiments provided a convenient way to assess proteome-wide glycosylation, but mechanistic interpretation is challenging. Adaptors, or proteins that recruit substrates to OGT’s active site, are proposed to play a substantial role in substrate selection.^*33-38*^ Thus, changes in global glycosylation with the TPR mutants could reflect changes in adaptor recognition rather than intrinsic differences in substrate recognition. To assess how individual substrates are affected by mutations in different regions of the TPR domain, we tested three previously studied OGT substrates, TAB1, CAMKIV, and CARM1.^*33, 39-42*^ These proteins are among the few substrates that have been characterized kinetically,^*43*^ and each is approximately 55 kDa with an enzymatic domain and a long, C-terminal region that is predicted to be disordered. Each also has one major glycosite that is located in a different region of the protein, either in a long loop in the folded domain (CAMKIV), in the unstructured region but immediately adjacent to the folded domain (TAB1), or in the unstructured region close to the C-terminus of the protein (CARM1) (Figure 4A). To identify large differences in how changing OGT’s TPR domain affects these substrates, we ran glycosylation assays under forcing conditions that would obscure small effects.

**Figure 4.**
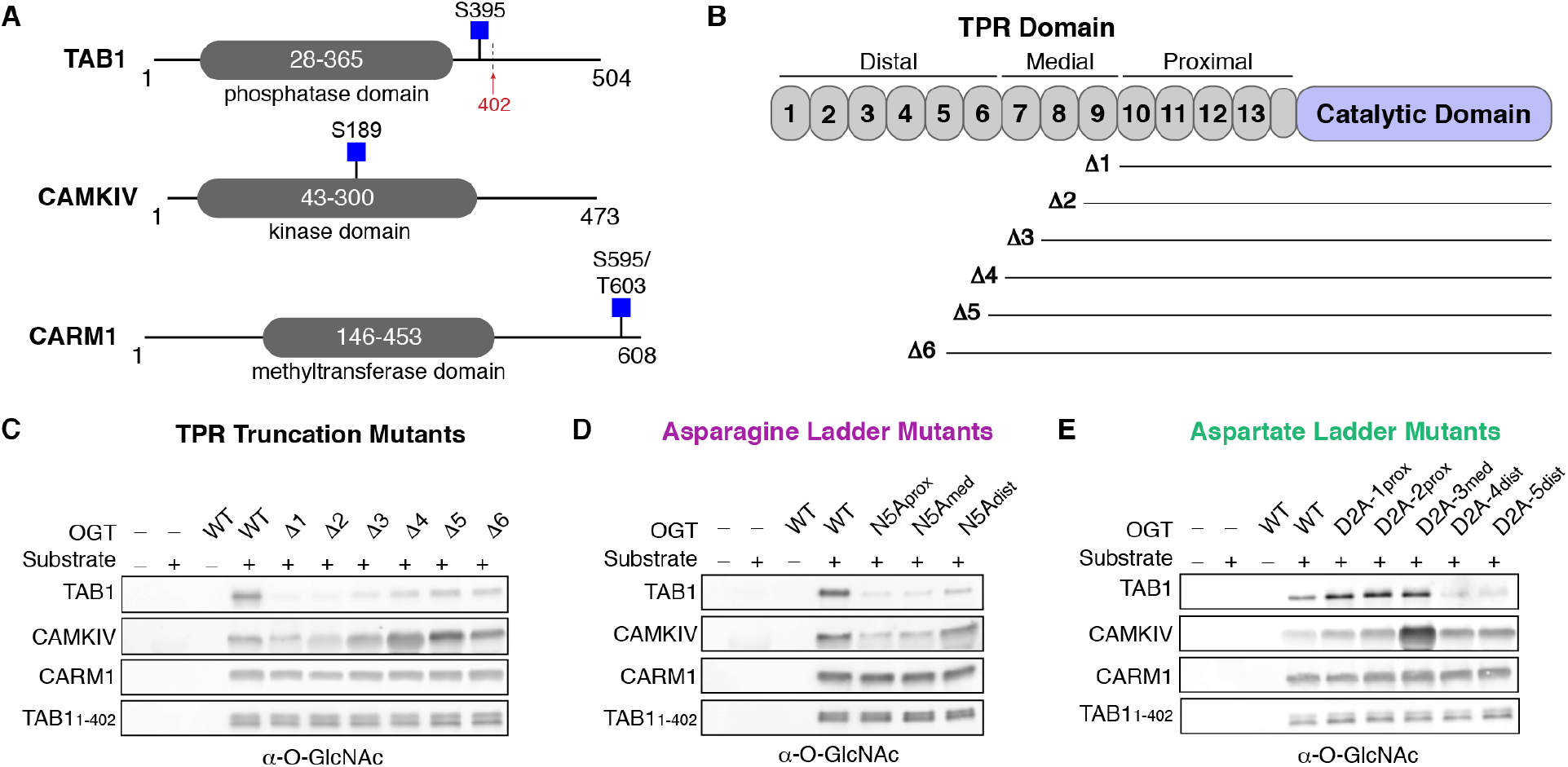
OGT protein substrates have different dependencies on the TPR domain for glycosylation. (A) Cartoon schematics of TAB1, CAMKIV, and CARM1 protein architecture. Glycosites are denoted by blue squares. TAB1 is proteolytically cleaved at residue K402 (red arrow). (B) Cartoon schematic of OGT N-terminal TPR truncation constructs. (C) *In vitro* glycosylation of substrates by TPR truncation mutants. (D) *In vitro* glycosylation of substrates by asparagine ladder mutants. (E) *In vitro* glycosylation of substrates by aspartate ladder mutants. Representative blots are shown for two independent experiments for each set of mutants.

We first assessed how truncating the TPR domain affected glycosylation of these three substrates (Figure 4B). Previous studies of two purified proteins showed that glycosylation was largely abolished when three N-terminal TPRs were removed, while glycosylation of a short peptide was unaffected.^*44, 45*^ These studies were limited to two substrates, but it has largely been assumed that all protein substrates must interact with a large part of the TPR domain to be glycosylated. We found that TAB1 behaved like the proteins in earlier studies in that removal of four or more TPRs greatly attenuated glycosylation (Figure 4C). In contrast, removing four to six N-terminal TPRs *increased* glycosylation of CAMKIV (Figures 4B,C: compare Δ6, which has 9 TPRs, to Δ4 which has 7 TPRs), although further TPR trimming led to reduced glycosylation levels compared to wildtype OGT. For CARM1, removal of as many as nine TPRs had no impact on glycosylation. Similar to CARM1, a TAB1 proteolysis product that co-purified with full-length TAB1^*19, 43*^ (TAB1_1-402_; Figure 4A) was glycosylated to the same extent by all truncation mutants. The results for CARM1 and TAB1_1-402_ show that proteins with glycosites very close to the C-terminus behave like peptide substrates in that they do not require contacts to the TPR domain to access the OGT active site for glycosylation.^*17, 44-46*^ Overall, the data for the three full-length protein substrates showed that each relies on contacts to different regions of OGT.

Although the truncation studies revealed differences in how proteins interact with the TPR domain, they did not provide information on whether contacts to the TPR lumen were important. We used the asparagine ladder mutants to assess whether substrates bind in the TPR lumen (Figure 2A,4D). As expected from the truncation studies, CARM1 and TAB1_1-402_ glycosylation was unaffected by the lumenal mutations. However, glycosylation of full-length TAB1 was largely abolished for all three of the N5A mutants. CAMKIV glycosylation was reduced for the N5A_prox_ and N5A_med_ mutants, but not for N5A_dist_. Therefore, both TAB1 and CAMKIV engage the TPR lumen, but TAB1 requires asparagine ladder residues in the distal region of the TPR lumen for glycosylation, whereas CAMKIV glycosylation depends only on contacts to the proximal and medial regions of the TPR lumen.

TAB1 and CAMKIV, the two substrates that interacted with the TPR lumen, also showed notable differences in glycosylation with the aspartate mutants (Figure 3A,4E). Whereas glycosylation of full-length TAB1 decreased when aspartates in the distal TPR lumen were mutated to alanine (D2A-4_dist_ – D2A-5_dist_), it increased for D2A mutants elsewhere (D2A-1_prox_ – D2A-3_med_). CAMKIV showed a large increase in glycosylation when two aspartates in the medial lumen were changed to alanines, but the other aspartate mutants showed only modest increases compared to wildtype OGT. Taken together, these findings are generally consistent with the proteome-wide studies in showing that replacing aspartates in the proximal and medial TPR lumen to alanine increases glycosylation. The appearance of new product bands in the proteome-wide studies indicated that new substrates were glycosylated, consistent with our previous findings that the aspartates can act as gatekeeper residues to limit protein glycosylation.^*23*^ The increased glycosylation of individual substates suggested that substrate turnover may also increase when the aspartates are removed. Consistent with this, we quantified turnover for TAB1 using wildtype OGT and the D2A-2_prox_ and found a 10-fold increase in turnover for the D2A mutant (Figure S2).

Our findings show that the conserved asparagine and aspartate ladders that span the TPR lumen regulate glycosylation of a large fraction of OGT’s glycoproteome, and also reveal major differences in how different substrates engage the lumen. The three purified substrates tested here provide different working models for substrate engagement during glycosylation. In one model (CARM1-like), substrates bind similarly to short peptides in the sense that they can access the active site without engaging the TPR lumen. In a second model (TAB1-like) substrates simultaneously engage the active site and the full TPR lumen to achieve glycosylation. The results for TAB1 are remarkable because they indicate an interaction with OGT that spans as much as 120 Å. Because removal of most of the C-terminal disordered domain removed TAB1’s dependence on lumenal contacts for glycosylation, we infer that the presence of this disordered domain somehow interferes with glycosylation. One function of the TPR domain may be to chaperone unfolding of the disordered domain so that the glycosylation site is accessible. In other contexts, TPR domain-containing proteins are known to serve as co-chaperones to regulate protein fate.^*47-50*^ Given our finding that substrate proteolysis can alter requirements for glycosylation, it is also possible that proteolysis serves to regulate some O-GlcNAc modifications.

The CAMKIV results support a third model for TPR binding in which substrates rely on contacts to only part of the TPR lumen. Unlike the other model substrates tested, the glycosite of CAMKIV is located in a long, disordered loop within the kinase domain. The location of the glycosite in a long loop may place topological constraints on how the substrate engages the TPR lumen^*42*^, and these constraints could explain the observation that removing distal TPRs increases glycosylation. The CAMKIV loop also contains a regulatory phosphorylation site, T200, that blocks its glycosylation.^*42, 43*^ This phosphorylation site is 11 residues C-terminal to the glycosite. Our earlier studies showed a strong bias against negatively-charged side chains several residues C-terminal to the site of glycosylation in OGT substrates, and we interpreted this bias as reflecting unfavorable interactions between substrate side chains and lumenal aspartates.^*23*^ We speculate that phosphorylation may prevent engagement of CAMKIV with OGT’s TPR lumen by introducing a repulsive interaction, providing one possible mechanism to explain “crosstalk” between phosphorylation and glycosylation.^*51*^

Our work, along with a recent study on how glycosylation of O-GlcNAcase (OGA) is affected when lumenal residues in OGT’s TPR domain are altered, is beginning to clarify how different substrates use OGT’s TPR domain to access its active site, pointing the way for future work to develop a molecular understanding of substrate recognition. The work on OGA showed that it behaves like TAB1 in using lumenal interactions that involve most of the TPR domain for substrate recognition.^*21, 52, 53*^ Further progress in understanding OGT substrate recognition will require integration of glycosylation assays with quantitative assays to measure substrate binding affinities, which so far do not exist, and ideally with structural information. In the meantime, we note that modulating glycosylation of broad subsets of substrates by changing features of the TPR domain may help clarify the function of O-GlcNAcylation in heath and disease.

## ASSOCIATED CONTENT

### Supporting Information

The following files are available free of charge.

Experimental protocols used in this study, supplemental figures S1-S5, and tables S1-S6 (PDF)

## AUTHOR INFORMATION

**Corresponding Author**

*Suzanne Walker – Department of Microbiology, Harvard Medical School, 4 Blackfan Circle, Boston, MA, 02115, USA;

*Cassandra M. Joiner – Department of Chemistry, St. Olaf College, 1520 St. Olaf Ave., Northfield, MN 55057, USA

**Present Addresses**

† Cassandra M Joiner – Department of Chemistry, St. Olaf College, 1520 St. Olaf Ave., Northfield, MN 55057, USA

† John Janetzko – Department of Molecular and Cellular Physiology, Stanford University School of Medicine, 279 Campus Drive, Stanford, CA 94305, USA

### Notes

The authors declare no competing financial interest.

## ACKNOWLEDGMENT

We thank Dr. David Vocadlo for providing the pET28a-TAB1, pET28a-CARM1, and pET28a-CAMKIV plasmids.

## TOC Figure

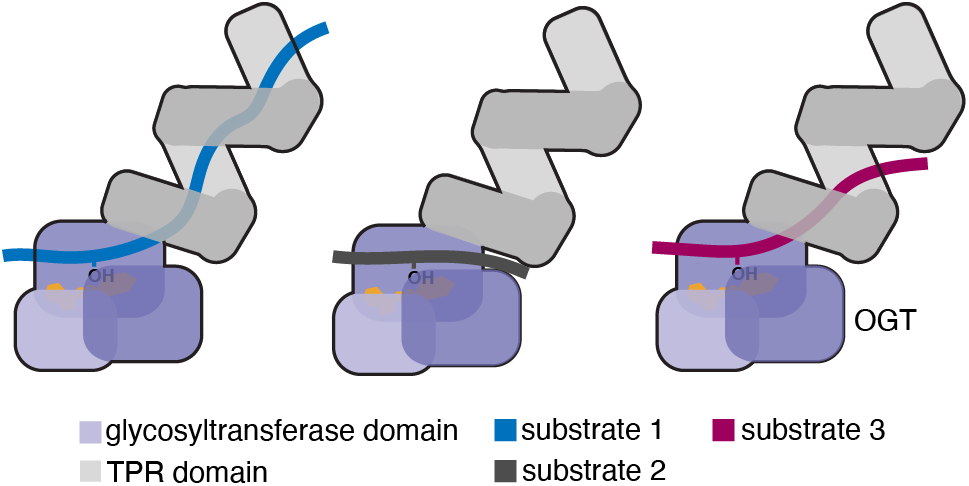

