## Supplemental Information for "Protein substrates engage the lumen of O-GlcNac transferase’s tetratricopeptide repeat domain in different ways"

| | |
|---|---|
| Supplemental Table S1-S2 | S2 |
| Supplemental Table S3-S5 | S3 |
| Supplemental Figures | S4 |
| Supplemental Table S6 | S5 |
| General Methods | S9 |
| Tables of Plasmids | S9 |
| Tables of Primers | S9 |
| Table of Gene Blocks | S10 |
| Construction of pET24-8XHIS-HRV3C-ncOGT variants | S10 |
| Bacterial expression and purification of pET24-8XHIS-HRV3C-ncOGT variants | S10 |
| *In vitro* glycosylation of HeLa extracts by ncOGT variants | S11 |
| Bacterial expression and purification of OGT substrates | S11 |
| *In vitro* glycosylation of OGT substrates by ncOGT variants | S12 |
| *In vitro* turnover assay of TAB1 glycosylation by ncOGT variants | S12 |
| References | S12 |

**Supplemental Table S1. OGT Asparagine Ladder Conservation Table**

| | % Conservation per amino acid | |
|---|---|---|
| **Residue** | **N** | **Next highest cons. residues** |
| N458 | 88.9 | 2.7 - S |
| N424 | 96.3 | 1.0 – S, 1.0 - A |
| N390 | 95.6 | 1.7 - S |
| N356 | 95.9 | 1.7 - S |
| N322 | 95.9 | 1.0 - D |
| N288 | 94.2 | 1.7 - D |
| N254 | 91.8 | 1.7 - S, 1.7 - K |
| N220 | 90.7 | 3.2 - S |
| N186 | 93.1 | 3.0 - A |
| N155 | 82.7 | 4.1 - D |
| N118 | 83.6 | 7.6 - L |
| N84 | 87.6 | 3.6 - S |

Conservation scores determined for OGT's TPR domain using the 1W3B PDB file and ConSurf Server.[1-3] 150 sequence homologs were compared.

**Supplemental Table S2. OGT Aspartate Ladder Conservation Table**

| | % Conservation per amino acid | |
|---|---|---|
| **Residue** | **D** | **Next highest cons. residues** |
| D454 | 54.5 | 33.8 - E |
| D420 | 53.2 | 38.1 - E |
| D386 | 81.8 | 9.8 - E |
| D318 | 49.5 | 11.9 - E |
| D284 | 55.9 | 34.8 - E |
| D216 | 80.6 | 13.6 - E |
| D152* | 45.9 | 48.1 - N |
| D114 | 85.8 | 4.4 - E |

* Position 152 can be either an aspartate (D – 45.9% conservation) or asparagine (N- 48.1% conservation). Conservation scores determined for OGT's TPR domain using the 1W3B PDB file and ConSurf Server.[1-3] 150 sequence homologs were compared.

**Supplemental Table S3. Quantification of *in vitro* glycosylation of purified substrates by OGT TPR truncation mutants** (average of 2 replicates)

| **OGT constructs** | **TAB1 (1-504) Glycosylation Signal** (Fold change compared to OGT WT) | **TAB1 (1-402) Glycosylation Signal** (Fold change compared to OGT WT) | **CARM1 Glycosylation Signal** (Fold change compared to OGT WT) | **CAMKIV Glycosylation Signal** (Fold change compared to OGT WT) |
|---|---|---|---|---|
| OGT WT | 1.0 | 1.0 | 1.0 | 1.0 |
| OGT Δ1 | 0.07 | 0.9 | 0.8 | 0.4 |
| OGT Δ2 | 0.05 | 0.8 | 0.7 | 0.5 |
| OGT Δ3 | 0.1 | 1.7 | 0.8 | 1.4 |
| OGT Δ4 | 0.2 | 1.6 | 0.8 | 2.0 |
| OGT Δ5 | 0.3 | 1.6 | 0.9 | 2.5 |
| OGT Δ6 | 0.2 | 1.7 | 1.0 | 1.7 |

**Supplemental Table S4. Quantification of *in vitro* glycosylation of purified substrates by OGT N5A mutants** (average of 2 replicates)

| **OGT constructs** | **TAB1 (1-504) Glycosylation Signal** (Fold change compared to OGT WT) | **TAB1 (1-402) Glycosylation Signal** (Fold change compared to OGT WT) | **CARM1 Glycosylation Signal** (Fold change compared to OGT WT) | **CAMKIV Glycosylation Signal** (Fold change compared to OGT WT) |
|---|---|---|---|---|
| OGT WT | 1.0 | 1.0 | 1.0 | 1.0 |
| OGT $N5A_{prox}$ | 0.1 | 1.0 | 1.0 | 0.4 |
| OGT $N5A_{med}$ | 0.1 | 1.0 | 1.0 | 0.5 |
| OGT $N5A_{dist}$ | 0.2 | 1.0 | 0.90 | 1.1 |

**Supplemental Table S5. Quantification of *in vitro* glycosylation of purified substrates by OGT D2A mutants** (average of 2 replicates)

| **OGT constructs** | **TAB1 (1-504) Glycosylation Signal** (Fold change compared to OGT WT) | **TAB1 (1-402) Glycosylation Signal** (Fold change compared to OGT WT) | **CARM1 Glycosylation Signal** (Fold change compared to OGT WT) | **CAMKIV Glycosylation Signal** (Fold change compared to OGT WT) |
|---|---|---|---|---|
| OGT WT | 1.0 | 1.0 | 1.0 | 1.0 |
| OGT D2A-$1_{prox}$ | 2.3 | 1.1 | 1.0 | 1.5 |
| OGT D2A-$2_{prox}$ | 2.3 | 1.1 | 1.3 | 2.0 |
| OGT D2A-$3_{med}$ | 2.2 | 1.0 | 1.4 | 5.3 |
| OGT D2A-$4_{dist}$ | 0.9 | 1.0 | 1.2 | 2.3 |
| OGT D2A-$5_{dist}$ | 0.7 | 1.2 | 1.5 | 2.3 |

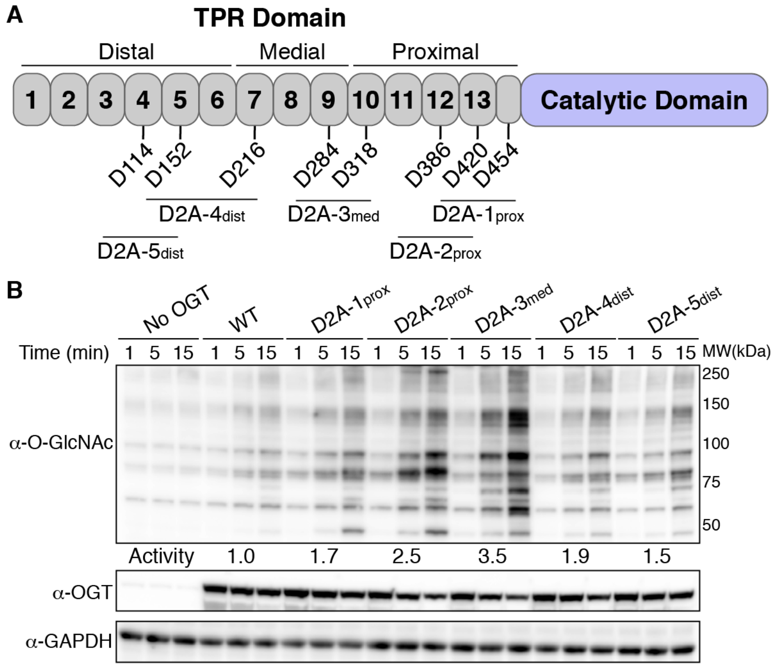


**Supplemental Figure S1. Aspartate ladder differentially regulates proteome-wide glycosylation.** (A) Cartoon schematic of OGT D2A mutant constructs. (B) Glycosylation of HeLa extracts show differential increases in glycosylation for D2A mutants with largest increases for D2A-2 and D2A-3. Representative blot for two independent experiments. See methods for relative glycosylation activity calculations.

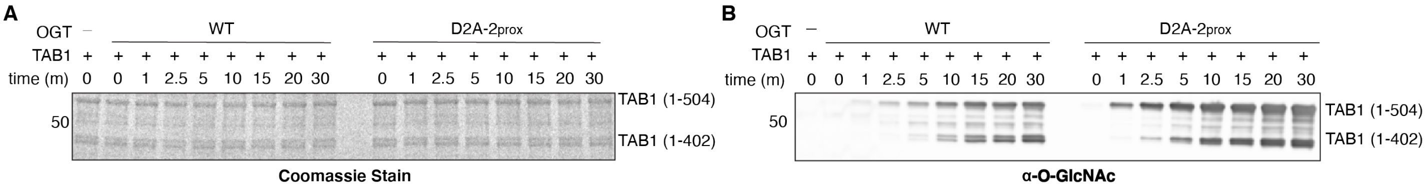


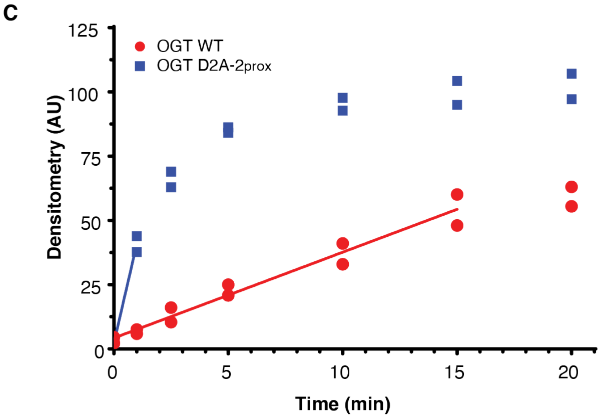


D

| OGT variant | TAB1 turnover | Fold change (over WT) |
|---|---|---|
| OGT WT | 3.3 ± 0.2 | 1.0 |
| OGT D2A-2prox | 39 ± 3 | 11.6 |

**Supplemental Figure S2. *In vitro* turnover of TAB1 by OGT WT and OGT D2A-2$_{prox}$.** A) Full-length SDS-PAGE gel and western blot showing time-dependence of *in vitro* glycosylation of TAB1 by OGT WT and OGT D2A-2$_{prox}$. A representative blot of two biological replicates shown. All SDS-PAGE gels were stained with Pierce Imperial$^{TM}$ protein stain and western blots were visualized using the pan O-GlcNAc antibody CTD110.6. B) Densitometry vs. time plot of *in vitro* TAB1 glycosylation by OGT WT and OGT D2A-2$_{prox}$. Best-fit lines are shown in the linear ranges of each activity curve. C) Comparison of TAB1 turnover calculated from linear ranges of glycosylation activity curves.

Supplemental Table S6. **Quantification of *in vitro turnover* of TAB 1 glycosylation by OGT WT and D2A-2$_{prox}$**

| Time (min) | OGT WT_Rep 1 (densitometry) | OGT WT_Rep 2 (densitometry) | OGT D2A-2prox_Rep1 (densitometry) | OGT D2A-2prox_Rep2 (densitometry) |
|---|---|---|---|---|
| 0.0 | 2.234 | 4.543 | 2.253 | 2.15 |
| 1.0 | 7.543 | 5.873 | 43.834 | 37.704 |
| 2.5 | 16.053 | 10.394 | 68.946 | 62.87 |
| 5.0 | 24.994 | 20.803 | 86.191 | 84.066 |
| 10.0 | 41.023 | 32.947 | 92.722 | 97.689 |
| 15.0 | 60.084 | 47.989 | 94.988 | 104.261 |
| 20.0 | 63.038 | 55.494 | 97.128 | 107.027 |
| 30.0 | 62.135 | 64.208 | 97.939 | 98.096 |

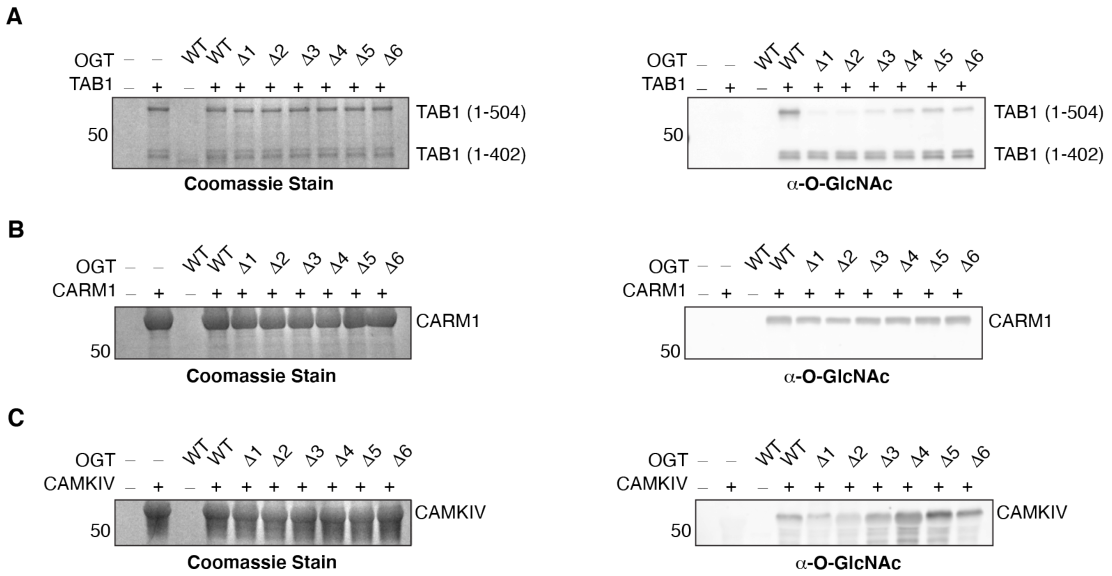


**Supplemental Figure S3. SDS-PAGE gels and western blots of *in vitro* glycosylation of OGT substrates by OGT TPR truncation variants.** A) *In vitro* glycosylation of TAB1 by 5N5A mutants. B) *In vitro* glycosylation of CARM1 by 5N5A mutants. C) *In vitro* glycosylation of CAMKIV by 5N5A mutants. All SDS-PAGE gels were stained with Instant Blue Coomassie stain and western blots were visualized using the pan O-GlcNAc antibody CTD110.6.

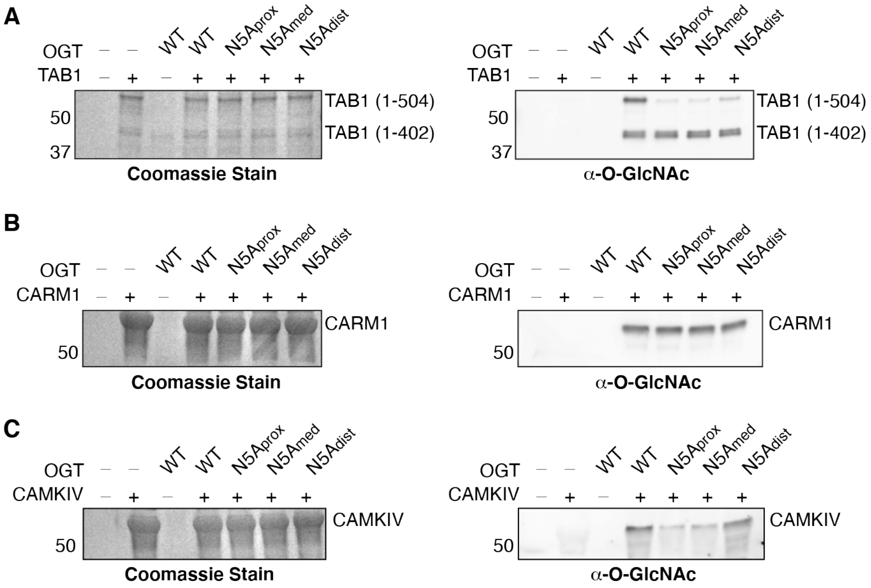


**Supplemental Figure S4. SDS-PAGE gels and western blots of *in vitro* glycosylation of OGT substrates by OGT asparagine ladder variants.** A) *In vitro* glycosylation of TAB1 by N5A mutants. B) *In vitro* glycosylation of CARM1 by N5A mutants. C) *In vitro* glycosylation of CAMKIV by N5A mutants. All SDS-PAGE gels were stained with Instant Blue Coomassie stain and western blots were visualized using the pan O-GlcNAc antibody CTD110.6.

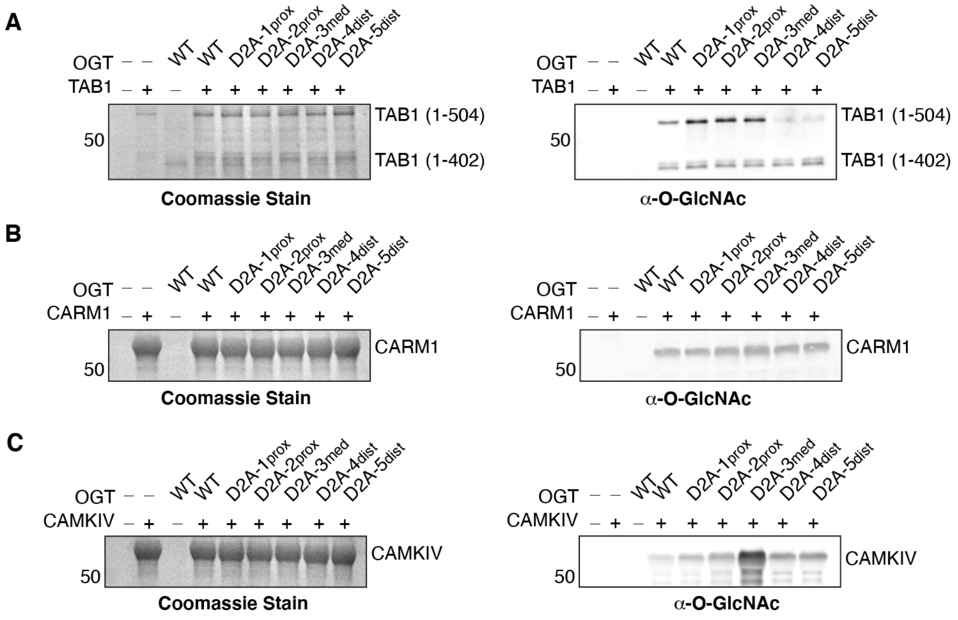


**Supplemental Figure S5. Full length SDS-PAGE gels and western blots of *in vitro* glycosylation of OGT substrates by OGT aspartate ladder variants.** A) *In vitro* glycosylation of TAB1 by D2A mutants. B) *In vitro* glycosylation of CARM1 by D2A mutants. C) *In vitro* glycosylation of CAMKIV by D2A mutants. All SDS-PAGE gels were stained with Instant Blue Coomassie stain and western blots were visualized using the pan O-GlcNAc antibody CTD110.6.

**General Methods**
All primers and gene blocks were purchased from Integrated DNA Technologies (IDT) and all isolated plasmid were verified by sequencing by the Dana-Farber/Harvard Cancer Center DNA Resource Core (Boston, MA).

**Table of Plasmids**

| Plasmids | Function | Reference |
|---|---|---|
| pET24b-8XHIS-HRV3C-ncOGT WT | Expresses 8XHIS-HRV3C-ncOGT (1-1036) (Uniport Accession: O15294) | 6 |
| pET24b-8XHIS-HRV3C-ncOGT N5A$_{prox}$ | Expresses 8XHIS-HRV3C-ncOGT (1-1036) with GCA codon replacing N322, N356, N390, N424, and N458 | 7,8 |
| pET24b-8XHIS-HRV3C-ncOGT N5A$_{med}$ | Expresses 8XHIS-HRV3C-ncOGT (1-1036) with GCA codon replacing N186, N220, N254, N288, and N322 | This study |
| pET24b-8XHIS-HRV3C-ncOGT N5A$_{dist}$ | Expresses 8XHIS-HRV3C-ncOGT (1-1036) with GCA codon replacing N84, N118, N155, N186, and N220 | This study |
| pET24b-8XHIS-HRV3C-ncOGT D420A/D454A (D2A-1$_{prox}$) | Expresses 8XHIS-HRV3C-ncOGT (1-1036) with GCA codon replacing the existing amino acids | 9 |
| pET24b-8XHIS-HRV3C-ncOGT D386A/D420A (D2A-2$_{prox}$) | Expresses 8XHIS-HRV3C-ncOGT (1-1036) with GCA codon replacing the existing amino acids | 9 |
| pET24b-8XHIS-HRV3C-ncOGT D284A/D318A (D2A-3$_{med}$) | Expresses 8XHIS-HRV3C-ncOGT (1-1036) with GCA codon replacing the existing amino acids | This study |
| pET24b-8XHIS-HRV3C-ncOGT D152A/D216A (D2A-4$_{dist}$) | Expresses 8XHIS-HRV3C-ncOGT (1-1036) with GCA codon replacing the existing amino acids | This study |
| pET24b-8XHIS-HRV3C-ncOGT D114A/D152A (D2A-5$_{dist}$) | Expresses 8XHIS-HRV3C-ncOGT (1-1036) with GCA codon replacing the existing amino acids | This study |
| pET24b-8XHIS-HRV3C-ncOGT D284A/D386A/D420A/D454A (D4A) | Expresses 8XHIS-HRV3C-ncOGT (1-1036) with GCA codon replacing the existing amino acids | This study |
| pET24b-8XHIS-HRV3C-ncOGT Δ1 | Expresses 8XHIS-HRV3C-ncOGT (323-1036) containing 4.5 TPRs (TPRs 10-13.5) | 5 |
| pET24b-8XHIS-HRV3C-ncOGT Δ2 | Expresses 8XHIS-HRV3C-ncOGT (289-1036) containing 5.5 TPRs (TPRs 9-13.5) | This study |
| pET24b-8XHIS-HRV3C-ncOGT Δ3 | Expresses 8XHIS-HRV3C-ncOGT (238-1036) containing 6.5 TPRs (TPRs 8-13.5) | This study |
| pET24b-8XHIS-HRV3C-ncOGT Δ4 | Expresses 8XHIS-HRV3C-ncOGT (221-1036) containing 7.5 TPRs (TPRs 7-13.5) | This study |
| pET24b-8XHIS-HRV3C-ncOGT Δ5 | Expresses 8XHIS-HRV3C-ncOGT (203-1036) containing 8.0 TPRs (TPRs 6.5-13.5) | This study |
| pET24b-8XHIS-HRV3C-ncOGT Δ6 | Expresses 8XHIS-HRV3C-ncOGT (169-1036) containing 9.0 TPRs (TPRs 5.5-13.5) | This study |
| pET28a-CARM1 | Expresses polyhistidine-tagged CARM1 (Uniport Accession: Q86X55) | 4 |
| pET28a-CAMKIV | Expresses polyhistidine-tagged CAMKIV (Uniport Accession: Q16566) | 4 |
| pET28a-TAB1 | Expresses polyhistidine-tagged TAB1 (Uniport Accession: Q15750) | 4 |

**Table of Primers**

| Primer ID | Sequence |
|---|---|
| ncOGT N5A-2 FOR | 5' - GCAGACCAAGCAACAGCGAAGTTCGGCTGGGTTTCG – 3' |
| ncOGT N5A-2 REV | 5' – GGCTAACATCAAACGTGAACAGGGTAACATCGAAGAAGCTG – 3' |
| ncOGT N5A-3 FOR | 5' – GAGTAAGCTTCAGCCAGCAGCGGGTTCTGTTTGATAGC – 3' |
| ncOGT N5A-3 REV | 5' – CACTGGGTAACGTTCTGAAAGAAGCTCGTATCTTCGACCGTGC – 3' |

| | |
|---|---|
| ncOGT D318A FOR | 5’ – GTGCCCGACCCACGCT**GCA**TCTCTGAACAACCTGG – 3’ |
| ncOGT D318A REV | 5’ - CCAGGTTGTTCAGAGA**TGC**AGCGTGGGTCGGGCAC – 3’ |
| ncOGT D284A FOR | 5’ - CAGCCGCACTTTCCA**GCA**GCTTACTGCAACCTGG – 3’ |
| ncOGT D284A REV | 5’ – CCAGGTTGCAGTAAGC**TGC**TGGAAAGTGCGGCTG – 3’ |
| ncOGT D216A FOR | 5’ – GGACCCGAACTTCCTG**GCA**GCTTACATCAACCTGG – 3’ |
| ncOGT D216A REV | 5’ – CCAGGTTGATGTAAGC**TGC**CAGGAAGTTCGGGTCC – 3’ |
| ncOGT D152A FOR | 5’ – CGTTCGTTCT**GCA**CTGGGTAACCTG – 3’ |
| ncOGT D152A REV | 5’ – CAGTACAGGTCCGGGTTG – 3’ |
| ncOGT D114A FOR | 5’ – GAAAACGGACTTCATC**GCA**GGTTACATCAACCTGGCTG – 3’ |
| ncOGT D114A REV | 5’ – CAGCCAGGTTGATGTAACC**TGC**GATGAAGTCCGGTTTC – 3’ |

**Table of Gene Blocks**

| Gene Block ID | Sequence |
|---|---|
| ncOGT N5A-2<br>(N186A/N220A/N254A/N288A<br>/N322A) gene block | 5’ – CGCTGTTGCTTGGTCT**GCA**CTGGGTTGCGTTTTCAACGCTCAGGGTGAA<br>ATCTGGCTGGCTATCCACCACTTCGAAAAAGCTGTTACCCTGGACCCGAACT<br>TCCTGGACGCTTACATC**GCA**CTGGGTAACGTTCTGAAAGAAGCTCGTCGTAT<br>CTTCGACCGTGCTGTTGCTGCTTACCTGCGTGCTCTGTCTCTGTCTCCGAAC<br>CACGCTGTTGTTCACGGT**GCA**CTGGCTTGCGTTTACTACGAACAGGGTCTGA<br>TCGACCTGGCTATCGACACCTACCGTCGTGCTATCGAACTGCAGCCGCACTT<br>TCCAGATGCTTACTGC**GCA**CTGGCTAACGCTCTGAAAAGAAAAAGGTTCTGTT<br>GCTGAAGCTGAAGACTGCTACAACACCGCTCTGCGTCTGTGCCCGACCCAC<br>GCTGACTCTCTGAAC**GCA**CTGGCTAACATCAAACGTGAAC – 3’ |
| ncOGT N5A-3<br>(N84A/N118A/N155A/N186A/<br>N220A)<br>gene block | 5’ – GCTGGCTGAAGCTTACTCT**GCA**CTGGGTAACGTTTACAAAGAACG<br>TGGTCAGCTGCAGGAAGCTATCGAACACTACCGTCACGCTCGCGTCT<br>GAAACCGGACTTCATCGACGGTTACATC**GCA**CTGGCTGCTGCTCTGG<br>TTGCTGCTGGTGACATGGAAGGTGCTGTTCAGGCTTACGTTTCTGCTC<br>TGCAGTACAACCCGGACCTGTACTGCGTTCGTTCTGACCTGGGT**GCA**<br>CTGCTGAAAGCTCTGGGTCGTCTGGAAGAAGCTAAAGCTTGCTACCT<br>GAAAGCTATCGAAACCCAGCCGAACTTCGCTGTTGCTTGGTCT**GCA**C<br>TGGGTTGCGTTTTCAACGCTCAGGGTGAAATCTGGCTGGCTATCCAC<br>CACTTCGAAAAAGCTGTTACCCTGGACCCGAACTTCCTGGACGCTTAC<br>ATC**GCA**CTGGGTAACGTTCTGAAAG – 3’ |

**Construction of pet24-8XHIS-HRV3C-ncOGT variants**

All plasmids containing mutations within ncOGT’s TPR domain were derived from the pET24b-8XHIS-HRV3C-ncOGT WT parent plasmid using the following methods:

<u>Asparagine ladder variants:</u> Using Gibson Assembly cloning, five sequential asparagine to alanine point mutations were made by inserting a ~450 base pair gene blocks containing the OGT sequences with the specific five sequence asparagine residues mutated to the GCA (alanine) codon. These gene blocks were designed to contain ~15-20 base pair of homology on either end to the site of incorporation in the pET24b-8XHIS-HRV3C-ncOGT WT plasmid. Primers were designed to linearize the pET24b-8XHIS-HRV3C-ncOGT WT plasmid at either side of the given gene block insertion. Each set of primers for a given 5N5A variant were used to amplify linearized version of the parent plasmid without the region containing the mutations. The amplified PCR products were gel extracted and incubated with the Gibson Assembly Master Mix (NEB) at 50 °C for 1 hour. The assembled plasmids were transformed into STELLAR cloning cells and confirmed by Sanger sequencing.

<u>Aspartate ladder and TPR truncation variants</u>: Using QuickChange site-directed mutagenesis, single point mutations were made by replacing the existing amino acid in the TPR lumen with the GCA (alanine) codon. PCR primers were designed to have ~15-20 base pairs of homology on either side of the mutation. Each set of primers for a given variant were used to amplify the TPR mutation using the parent plasmid as the template. The amplified PCR products were digested with the Dpn1 restriction enzyme to digest the parent plasmid, leaving the desired mutant plasmids. The digested DNA products were transformed into STELLAR or Nova Blue cloning cells and confirmed by Sanger sequencing.

**Bacterial expression and purification of 8XHIS-HRV3C-ncOGT variants**

Expression of 8XHIS-HRV3C-ncOGT WT, TPR truncation mutants, and TPR point mutants were carried out in LOBSTR (BL21-DE3) cells (Kerafast). 50 mL starter cultures of 50 μg/mL kanamycin supplemented LB media were grown at 37 ˚C (250 rpm) from a single colony before addition to 1.5 L of LB media supplemented with 50 μg/mL kanamycin. Cells were grown at 37 ˚C (250 rpm) to an OD600 of 1.0. The cells were cooled to 16 °C for 30 minutes, and expression was induced with 0.2 mM isopropyl β-D-1-thiogalactopyranoside (IPTG) overnight at 16 ˚C. The cells were pelleted by centrifugation at 4200 rpm for 20 minutes and stored at -80 ˚C until lysis.

For protein purifications, the cell pellets lysed and the purified using Ni-NTA agarose superflow resin (Qiagen) as previously described.[9] The eluates were supplemented with 1mM THP and concentrated using 100K MWCO Amicon concentration tubes (Millapore) for full-length ncOGT variants and 50K MWCO Amicon concentration tubes for the ncOGT TPR truncation variants. The proteins were isolated through size-exclusion chromatography using a Superdex 200 increase 10/300 GL column (GE Life Sciences) with 1X TBS, pH 7.4. The resulting fractions' identity were verified by SDS-PAGE with appropriate molecular weight standards. Fractions containing the ncOGT variants were pooled and concentrated using the 100K MWCO Amicon concentration tube (50K MWCO tubes for TPR truncations). The concentrated protein was aliquoted and stored at -80 ˚C until use. Before use, OGT aliquots were thawed on ice, and centrifuged at max speed for 20 minutes at 4 ˚C to pellet any aggregated protein. The supernatant was removed and store on ice until use, with the concentration measured using A280 with the extinction coefficient of 117580 $M^{-1}cm^{-1}$ for full-length ncOGT, 77240 $M^{-1}cm^{-1}$ for OGT 4.5, 80220 $M^{-1}cm^{-1}$ for OGT 5.5, 84690 $M^{-1}cm^{-1}$ for OGT 6.5, 82170 $M^{-1}cm^{-1}$ for OGT 7.5, 93170 $M^{-1}cm^{-1}$ for OGT 8.0, and 100160 $M^{-1}cm^{-1}$ for OGT 9.0.

***In vitro*** **glycosylation of HeLa extracts by ncOGT variants**

Protein glycosylation profiles for each ncOGT variants were determined in HeLa whole cell extracts that were prepared as previously described.[8] For these experiments, HeLa cell extracts were thawed on ice and centrifuged at 4 ˚C for 20 minutes at 45,000 rpm (Beckman Coulter TLA120.2 rotor) to pellet any remaining precipitated debris. Following centrifugation, the HeLa cell extracts were aliquoted to a final volume of 40 μL. The aliquots were incubated with 1 mM of UDP-GlcNAc and equal volumes of reaction buffer (20 mM Tris, pH 7.4, 150 mM NaCl, 20 mM MgCl2, 1 mM trishydroxypropylphosphine) and 1 μM OGT for ncOGT D2A and D4A mutants or 2.5 μM OGT for ncOGT N5A mutants at 37 ˚C. At 1 minute, 5 minutes, and 15 minutes, 10 μL aliquots were taken from the reaction and quenched with 10 μL 2X Laemmli loading buffer containing β-mercaptoethanol. The samples were boiled at 95 ˚C for 10 minutes and run on a 4-20% TGX SDS-PAGE gel (Bio-Rad) at 180 V for 1.5 hours. The gels were transferred to PVDF membranes (BioRad) and analyzed by western blot with the anti-O-GlcNAc (CTD110.6, Cell Signaling), anti-OGT (D1D8Q, Cell Signaling), and anti-GAPDH (9484, Abcam) antibodies. Activity (densitometry) calculated in ImageJ by measuring the entire lane for each mutant, normalizing to the corresponding WT lane, and using the following equation: $\frac{[\left(\frac{mut\ 1\ min}{wt\ 1\ min}\right)+\left(\frac{mut\ 5\ min}{wt\ 5\ min}\right)+\left(\frac{mut\ 15\ min}{wt\ 15\ min}\right)]}{3}$. Activity reported is an average of two biological replicates.

**Bacterial expression and purification of OGT substrates**

Expression of pET28a-CARM1, CAMKIV, and TAB1 were carried out in LOBSTR (BL21-DE3) cells (Kerafast). 50 mL starter cultures of 50 μg/mL kanamycin supplemented LB media were grown at 37 ˚C (250 rpm) from a single colony before addition to 1.5 L of LB media supplemented with 50 μg/mL kanamycin. Cells were grown at 37 ˚C (250 rpm) to an OD600 of 0.6. The cells were cooled to 16 °C for 30 minutes, and expression was induced with 0.2 mM isopropyl β-D-1-thiogalactopyranoside (IPTG) overnight at 16 ˚C. The cells were pelleted by centrifugation at 4200 rpm for 20 minutes and stored at -80 ˚C until lysis.

The cell pellets were resuspended in 1X TBS, pH 7.4 (50 mM Tris, pH 7.4, 150 mM NaCl, supplemented with 0.1 mg/mL lysozyme, 0.1 mg/mL DNAse I, and 1 mM PMSF) at 4 ˚C for 30 minutes. The cells were lysed using an Avestin Emulsiflex C3 cell disruptor (ATA Scientific) three times at 15,000 psi and the debris was pelleted through centrifugation at 14,000xg for 20 minutes at 4 ˚C. The supernatant was collected, and 40 mM imidazole was added before the lysate was incubated with Ni-NTA agarose superflow resin (Qiagen) for 1 hour at 4 ˚C, which was prewashed with 3 column volumes 1X TBS, pH 7.4 + 40 mM imidazole for nickel affinity purification. The flow through was removed and the resin was washed with 10 column volumes of 1X TBS, pH 7.4 + 50 mM imidazole. The protein was then eluted with 4 column volumes 1X TBS, pH 7.4 + 250 mM imidazole. The eluate supplemented with 1mM THP and concentrated using a 30K MWCO Amicon concentration tube (Millapore). The protein was isolated through size-exclusion chromatography using a Superdex 75 increase 10/300 GL column (GE Life Sciences) with 1X TBS, pH 7.4. The resulting fractions' identity were verified by SDS-PAGE with appropriate molecular weight standards. Fractions contains the ncOGT variants were pooled and concentrated using the 30K MWCO Amicon concentration tube. The concentrated protein was aliquoted and stored at -80 ˚C until use. Before use, OGT aliquots were thawed on ice, and centrifuged at max speed for 20 minutes at 4 ˚C to pellet any aggregated protein. The supernatant was removed and store on ice until use, with the concentration measured using A280 with the extinction coefficient of 61170 $M^{-1}cm^{-1}$ for CARM1, 48360 $M^{-1}cm^{-1}$ for CAMKIV, and 37360 $M^{-1}cm^{-1}$ for TAB1.

***In vitro* glycosylation of OGT substrates by ncOGT variants**
ncOGT variants and the purified substrates, CARM1, CAMKIV, and TAB1 were thawed on ice, and centrifuged at 4 ˚C for 20 minutes at max speed (Eppendorf 5424R) to pellet any remaining precipitated debris. Following centrifugation, 2.5 μM OGT, 15 μM substrate, 1 mM UDP-GlcNAc were incubated in reaction buffer (20 mM Tris, pH 7.4, 150 mM NaCl, 20 mM MgCl2, 1 mM trishydroxypropylphosphine) at a final volume of 20 μL for 90 minutes at 37 °C, 350 rpm. The reactions were quenched with 20 μL 2X Laemmli loading buffer containing β-mercaptoethanol. The samples were boiled at 95 ˚C for 10 minutes and run on a 4-20% TGX SDS-PAGE gel (Bio-Rad) at 180 V for 1.5 hours. The gels were transferred to nitrocellulose membranes and analyzed by western blot with the anti-O-GlcNAc (CTD110.6, Cell Signaling) primary antibody and anti-mouse IgG-680RD IRDye (LI-COR) secondary antibody. Activity (densitometry) reported in Tables S3-5 calculated in ImageJ by measuring the densitometry of substrate glycosylation band for each mutant, normalizing to the corresponding WT lane using the following equation: $\frac{Mutant\ activity}{WT\ activity}$. Activity reported is an average of two replicates.

***In vitro* turnover assay of TAB1 glycosylation by ncOGT variants**
Turnover rates of TAB1 glycosylation for ncOGT WT and D2A-$2_{prox}$ were determined using an *in vitro* glycosylation time course followed by western blotting. For each reaction, 1 µM ncOGT variant, 20 µM TAB1, and 1 mM UDP-GlcNAc were incubated in reaction buffer (20 mM Tris, pH 7.4, 150 mM NaCl, 20 mM $MgCl_2$, 1 mM trishydroxypropylphosphine) at a total volume of 90 µL. Reactions were incubated at 37 °C with 350 rpm shaking. At 0, 1, 2.5, 5, 10, 15, and 20 minutes, 10 µL of each reaction was removed and quenched with 10 µL 2X Laemmli loading buffer containing β-mercaptoethanol. The samples were boiled at 95 °C for 10 minutes. The samples were split between two 4-20% TGX SDS-PAGE gels (Bio-Rad) and ran at 180V for 1.5 hours. One gel was stained with Pierce Imperial™ protein stain according to the manufacturer's protocol, while the other was transferred to a nitrocellulose membrane and analyzed by western blot with the anti-O-GlcNAc (CTD110.6, Cell Signaling) primary antibody and anti-mouse IgG-680RD IRDye (LI-COR) secondary antibody. Activity (densitometry) reported in Figure S2 was calculated in ImageJ for each timepoint by measuring the densitometry of the full-length glycosylated substrate band. Substrate turnover by ncOGT variants reported in Table S6 was calculated by fitting the linear ranges of the activity curves and taking the slopes of the resultant lines of best fit. Relative turnover was calculated using the following equation: $\frac{slope\ of\ mutant\ turnover}{slope\ of\ WT\ turnover}$. Data reported is an average of two replicates.
